# Vulnerabilities in coronavirus glycan shields despite extensive glycosylation

**DOI:** 10.1101/2020.02.20.957472

**Authors:** Yasunori Watanabe, Zachary T. Berndsen, Jayna Raghwani, Gemma E. Seabright, Joel D. Allen, Jason S. McLellan, Ian A. Wilson, Thomas A. Bowden, Andrew B. Ward, Max Crispin

## Abstract

Severe acute respiratory syndrome (SARS) and Middle East respiratory syndrome (MERS) coronaviruses (CoVs) are zoonotic pathogens with high fatality rates and pandemic potential. Vaccine development has focussed on the principal target of the neutralizing humoral immune response, the spike (S) glycoprotein, which mediates receptor recognition and membrane fusion. Coronavirus S proteins are extensively glycosylated viral fusion proteins, encoding around 69-87 N-linked glycosylation sites per trimeric spike. Using a multifaceted structural approach, we reveal a specific area of high glycan density on MERS S that results in the formation of under-processed oligomannose-type glycan clusters, which was absent on SARS and HKU1 CoVs. We provide a comparison of the global glycan density of coronavirus spikes with other viral proteins including HIV-1 envelope, Lassa virus glycoprotein complex, and influenza hemagglutinin, where glycosylation plays a known role in shielding immunogenic epitopes. Consistent with the ability of the antibody-mediated immune response to effectively target and neutralize coronaviruses, we demonstrate that the glycans of coronavirus spikes are not able to form an efficacious high-density global shield to thwart the humoral immune response. Overall, our data reveal how differential organisation of viral glycosylation across class I viral fusion proteins influence not only individual glycan compositions but also the immunological pressure across the viral protein surface.

## Main

Coronaviruses (CoVs) are enveloped pathogens responsible for multiple respiratory disorders of varying severity in humans^1^. Certain CoVs represent a significant threat to global human health, as illustrated by outbreaks of severe acute respiratory syndrome coronavirus (SARS-CoV) in 2003^2^, Middle East respiratory syndrome coronavirus (MERS-CoV) in 2012^3^, and most recently of 2019-nCoV in Wuhan^4^. Given their mortality rates, the current lack of targeted treatments and licensed vaccines, and their capacity to transmit between humans and across species barriers^5–10^, there is an urgent need for effective countermeasures to combat these pathogens. Ongoing vaccine development efforts have primarily focused on the spike (S) proteins that protrude from the viral envelope and constitute the main target of neutralizing antibodies^11,12^.

These trimeric S proteins mediate host cell entry with the S1 and S2 subunits responsible for binding to the host cell receptor and facilitating membrane fusion, respectively^13–17^. MERS S binds to dipeptidyl-peptidase 4 (DPP4)^18^, whereas SARS S^19^ and 2019-nCoV^20^ utilize angiotensin-converting enzyme 2 (ACE2) as a host cellular receptor. CoV S proteins are the largest class I viral fusion proteins known^15^, and are extensively glycosylated, with SARS and MERS S glycoproteins both encoding 69 N-linked glycan sequons per trimeric spike with 2019-nCoV containing 66 sites. These extensive post-translational modifications often mask immunogenic protein epitopes from the host humoral immune system by occluding them with host-derived glycans^21,22^. This phenomenon of immune evasion by molecular mimicry and glycan shielding has been observed and well characterised across other viral glycoproteins, such as HIV-1 envelope protein (Env)^23–25^, influenza hemagglutinin (HA)^26–28^ and Lassa virus glycoprotein complex (LASV GPC)^29–31^.

Previous analyses of viral glycan shields have revealed the presence of underprocessed oligomannose-type glycans that seemingly arise due to steric constraints that prevent access of glycan processing enzymes to substrate glycans^29,32,33^, especially when the viral glycoprotein has evolved to mask immunogenic epitopes with a particularly dense array of host-derived glycans^31,34^. Restricted access to these glycan sites or interference with surrounding protein surface or neighbouring glycan residues can render glycan processing enzymes ineffective in specific regions^32,33,35^. Glycan processing on soluble glycoproteins has also been shown to be a strong reporter of native-like protein architecture and thus immunogen integrity^36–38^; and glycan processing on a successful immunogen candidate should therefore mimic, as closely as possible, the structural features observed on the native virus^39,40^.

Here, we provide global and site-specific analyses of N-linked glycosylation on soluble SARS, MERS and HKU1 CoV S glycoproteins and reveal extensive heterogeneity, ranging from oligomannose-type glycans to highly processed complex-type glycosylation. Mapping of these glycans onto the structures of the trimeric S proteins revealed that some of these glycans contribute to the formation of a cluster of oligomannose-type glycans at specific regions of high glycan density on MERS-CoV S. Molecular evolution analysis of SARS and MERS S genes also reveals a higher incidence of amino-acid diversity on the exposed surfaces of the S proteins that are not occluded by N-linked glycans. Additionally, we compare the structures of the respective glycan coats of SARS and HIV-1 envelope proteins using cryogenic electron microscopy (cryo-EM) and computational modelling, which delineate a sparse glycan shield exhibited on SARS S compared to other viral glycoproteins. We therefore undertook a comparative analysis of viral glycan shields from characterized class I fusion proteins to highlight how glycosylation density influences oligomannose-type glycan abundance, and the relationship between effective glycan shields and viral evasion ability. Together, these data underscore the importance of glycosylation in viral immune evasion.

## Results & Discussion

### Glycan Processing of Trimeric SARS and MERS Spike Glycoproteins

In order to generate a soluble mimic of the viral S proteins, we used the 2P stabilised native-like SARS and MERS S protein antigens, the design and structures of which have been described previously by Pallesen et al.^41^. SARS, MERS and HKU1 S genes encode a large number of N-linked glycan sequons 23, 23 and 29, respectively (Fig. 1A). We initially sought to quantitatively assess the composition of the carbohydrate structures displayed on the S glycoproteins. To this end, N-linked glycans were enzymatically released, fluorescently labelled, and subjected to hydrophilic interaction chromatography-ultra-performance liquid chromatography (HILIC-UPLC). Cleavage of the fluorescently labelled glycans by endoglycosidase H (Endo H) revealed a population (SARS 32.2%; MERS 33.8%, HKU1 25.0%) of under-processed oligomannose-type glycans (Fig. 1B). This observation of both complex and oligomannose-type glycans reveals that the majority of N-linked glycans are able to be processed, although there is limited processing at specific sites across the S proteins. It is also interesting to note that the distribution of oligomannose-type glycans was broad, with Man_5_GlcNAc_2_ to Man_9_GlcNAc_2_ glycans all present, without one particular dominant peak, as is the case for some viral glycoproteins, such as HIV-1 Env^36^. The proportion of oligomannose-type glycans on recombinant coronavirus S proteins is consistent with previous studies performed on virally derived MERS and SARS coronavirus S proteins^22,42^. Coronaviruses have been previously been reported to form virions by budding into the lumen of endoplasmic reticulum-Golgi intermediate compartments (ERGIC)^43–45^. Observations of hybrid- and complex-type glycans on virally derived material^22,42^ would, however, suggest that it is likely that coronavirus virions travel through the Golgi apparatus after virion formation in the ERGIC en route to the cell surface, thus supporting recombinant immunogens as models of viral glycoproteins.

**Fig. 1.**
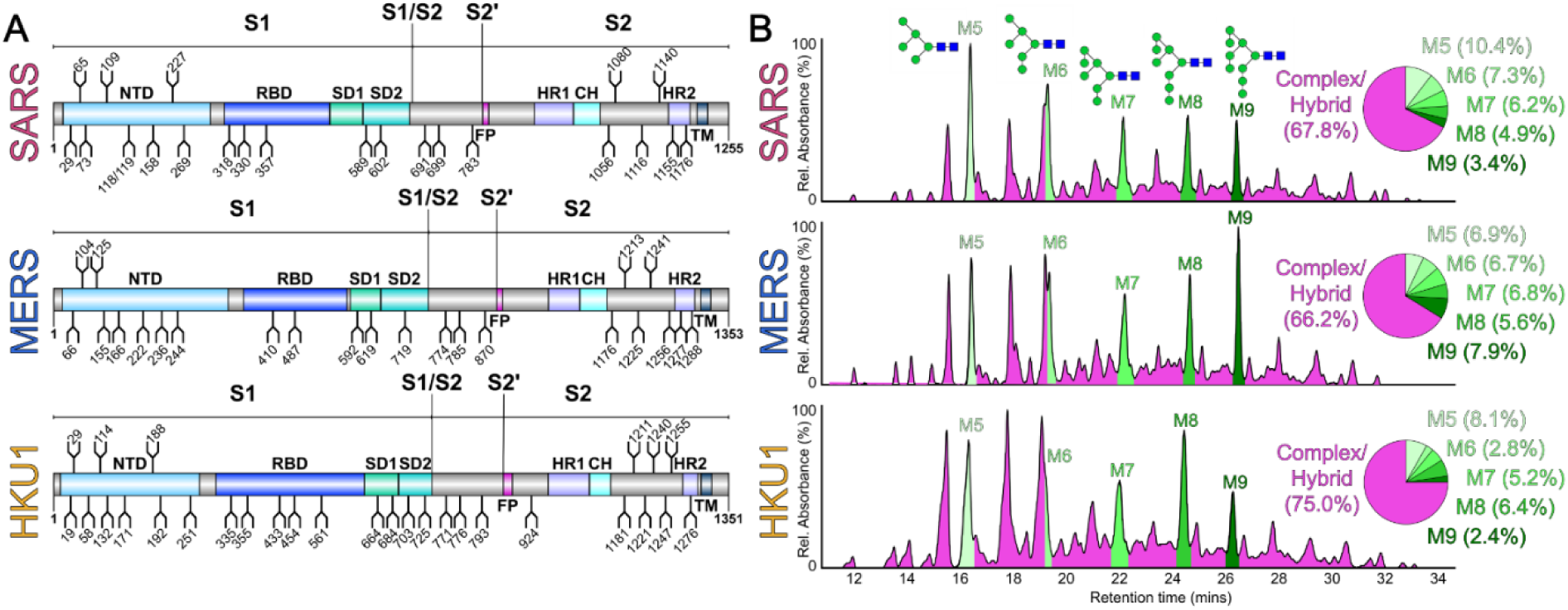
Compositional analysis of SARS, MERS and HKU1 glycans. **(A)** Schematic representation of SARS, MERS and HKU1 coronavirus S glycoproteins, showing the positions of N-linked glycosylation amino-acid sequons (NXS/T, where X ≠ P) shown as branches. The domains of the S glycoproteins are illustrated: N-terminal domain (NTD), receptor-binding domain (RBD), sub-domain 1/2 (SD1/2), fusion peptide (FP), heptad repeat 1/2 (HR1/2), central helix (CH), and transmembrane domain (TM). **(B)** HILIC-UPLC chromatograms of fluorescently labelled N-linked glycans from SARS, MERS and HKU1 S. Oligomannose-type glycans (M5 to M9; Man_5_GlcNAc_2_-Man_9_GlcNAc_2_) (green) and complex-type glycans (magenta) were identified by Endo H digestion, with quantification of major glycan types summarised as a pie chart. Oligomannose-type glycans are schematically annotated with mannose residues as green circles and GlcNAc residues as blue squares.

In order to ascertain the precise structures of N-linked glycans from the released pool, the glycans of each coronavirus S protein were analysed by negative-ion ion-mobility-electrospray ionisation mass spectrometry (IM-ESI MS) (Supplementary Fig. 1). Consistent with the UPLC data, IM-ESI MS confirmed an array of complex-type glycans ranging from mono-to tetra-antennary, but also oligomannose- and hybrid-type glycans. The glycan compositions characterised in the spectra were largely invariant among the coronaviruses with no major structural differences observed.

### Structural Clustering of Underprocessed Glycans on MERS S Head Region

Following UPLC and IM-ESI MS analysis of released N-linked glycans, we performed glycopeptide analysis to ascertain the compositions of glycans present at all of the potential N-linked glycosylation sites (PNGs). MERS, SARS and HKU1 recombinant S proteins were reduced, alkylated and digested with an assortment of proteases to yield glycopeptides, which were subjected to in-line liquid-chromatography mass spectrometry (LC-MS). This analysis revealed that each site presents differential levels of oligomannose, hybrid, and complex-type glycan populations (Fig. 2A & 2B). Using structures of the trimeric MERS and SARS S proteins (PDB ID: 5X59 and 5X58, respectively), we generated models of fully glycosylated coronavirus spikes using our experimentally determined glycan compositions (Fig. 3A and 3B). This analysis revealed that underprocessed oligomannose-type glycans on MERS S co-localize to a specific cluster on the head of the S protein, consisting of glycans at Asn155, Asn166, and Asn236 (Fig. 3A). We hypothesized that the fully oligomannose-type glycan population in this cluster arises due to the hindered accessibility of glycan processing enzymes to access the substrate glycan^33^. As such, we performed mutagenesis to knock out glycosylation sites with N155A, N166A, and N236A mutations. Site-specific analysis of these glycan-KO mutants revealed enhanced trimming of mannose residues, i.e. increased processing, when clustering of glycans was reduced (SI Fig. 4). The presence of clustered oligomannose-type glycans is reminiscent of that found on other viral glycoproteins, including HIV-1 Env and LASV GPC^29,36,46^.

**Fig. 2.**
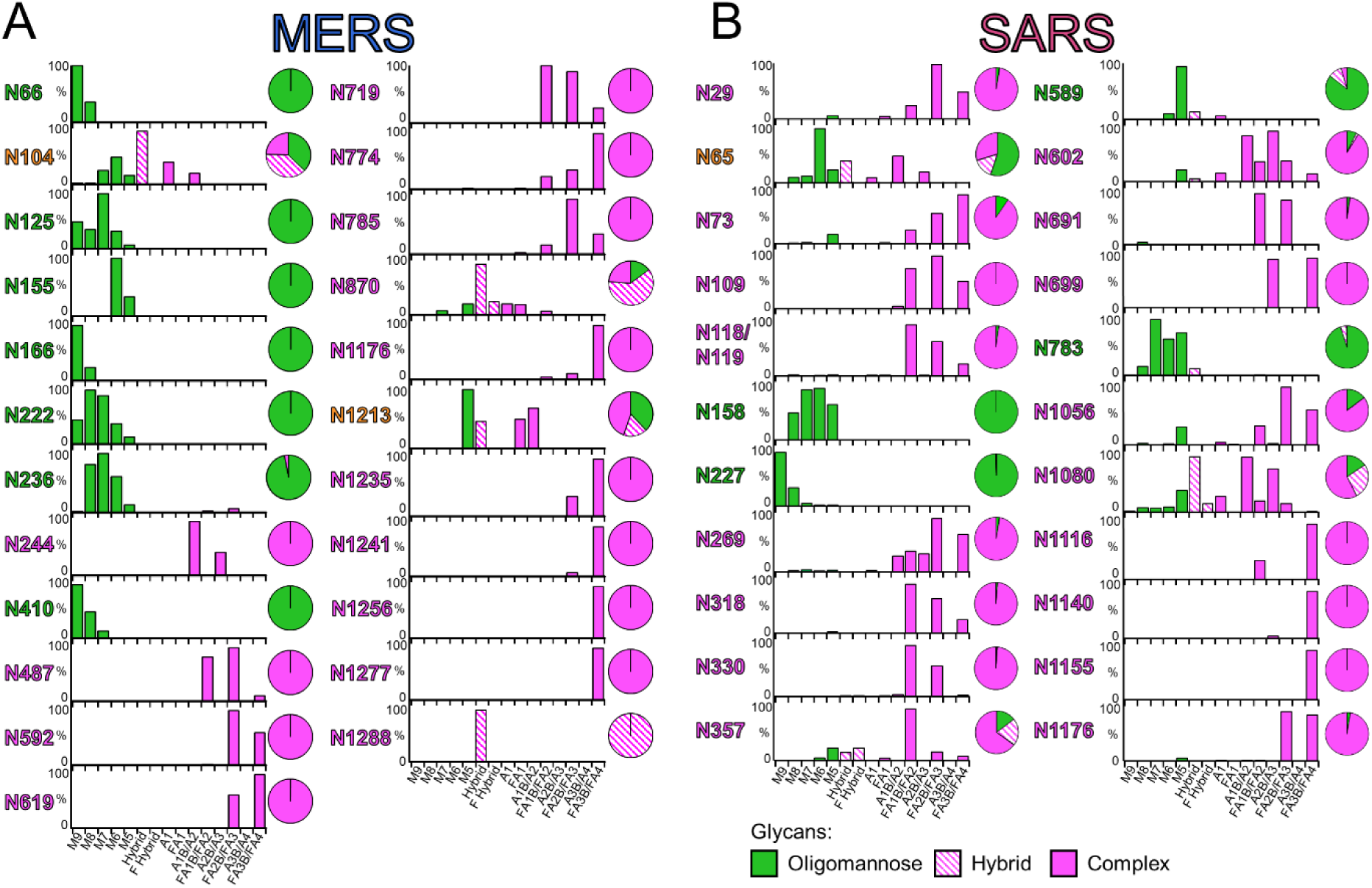
Quantitative site-specific N-linked glycan analysis of **(A)** MERS and **(B)** SARS S glycoproteins. Purified S proteins were digested with trypsin, chymotrypsin, alpha-lytic protease, Glu-C, and trypsin plus chymotrypsin, then analysed by LC-ESI MS. Glycan compositions are based on the glycan library generated from negative-ion mass spectrometry of released N-glycans. The bar graphs represent the relative quantities of each glycan group with oligomannose-type glycan series (M9 to M5; Man_9_GlcNAc_2_ to Man_5_GlcNAc_2_) (green), afucosylated and fucosylated hybrid glycans (Hybrid & F Hybrid) (dashed pink), and complex glycans grouped according to the number of antennae and presence of core fucosylation (A1 to FA4) (pink). Left to right; least processed to most processed. The pie charts summarise the quantification of these glycans. Additional compositional information regarding the distribution of fucosylation and sialylation can be found in SI Fig 3.

**Fig. 3.**
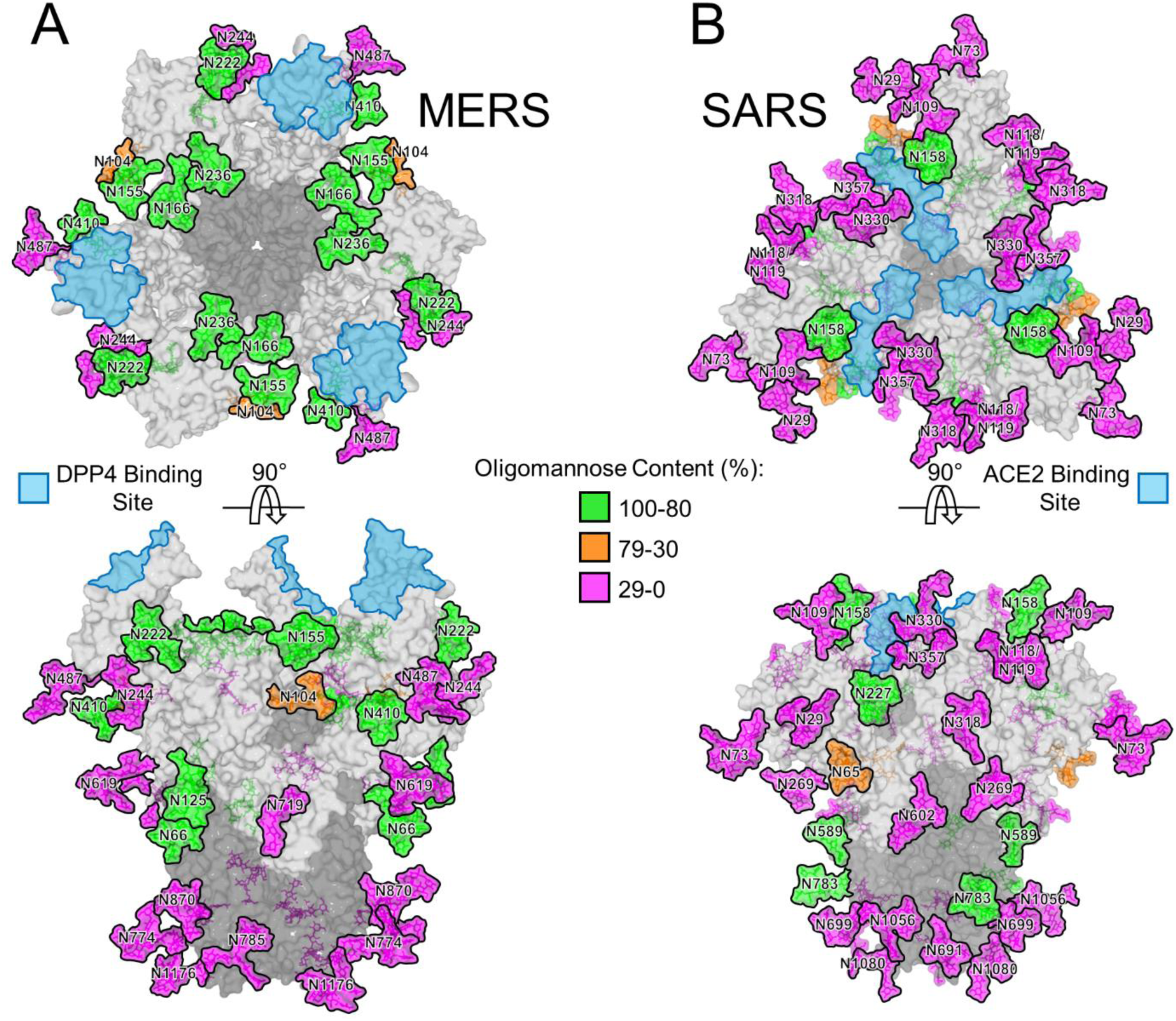
Structural-based mapping of N-linked glycans on **(A)** MERS and **(B)** SARS S proteins. The modelling of the experimentally observed glycosylation is illustrated on the pre-fusion structure of trimeric MERS S (PDB ID 5X59)^17^ and SARS S (PDB ID 5X58)^17^ glycoproteins. The glycans are colored according to oligomannose content, as defined by the key. DPP4 receptor binding sites and ACE2 receptor binding sites for MERS and SARS, respectively, are indicated in light blue. The S1 and S2 subunits are colored light grey and dark grey, respectively.

Interestingly, SARS and HKU1 (SI Fig. 2) S proteins did not exhibit a specific mannose cluster that contributes to the overall mannose abundance, but rather only isolated glycans were underprocessed. We would speculate that the oligomannose-type glycans here arise from protein-directed inhibition of glycan processing, as opposed to the glycan-influenced processing observed on MERS. The presence of oligomannose-type glycans has also been implicated in innate immune recognition of coronaviruses by lectins^47,48^ that recognise these underprocessed glycans as pathogen-associated molecular patterns.

Given that the receptor binding domain is the main target of neutralising antibodies^12^, it is surprising that the DPP4 receptor binding site of MERS S was not occluded at all by glycans (Fig. 3A), as observed for other receptor-binding sites of class I viral fusion proteins, including SARS S (Fig. 3B), HIV-1 Env^49^, LASV GPC^29^, and influenza HA^50,51^.

### Sequence Diversification Occurs at Antibody-accessible Regions of CoV spikes

We hypothesized that solvent-accessible, amino-acid residues on S proteins would be undergoing higher rates of mutations compared to buried residues and regions that are occluded by glycans, which are unable to be targeted by host immune responses. To that end, we performed an evaluation of amino-acid diversification on a residue-specific level, using publicly available gene sequences of SARS and MERS S. Firstly, we found that amino-acid diversity was elevated at known epitopes targeted by neutralizing antibodies, such as the N-terminal domain and the receptor binding domains, and reduced in the regions in the S2 domain, such as the fusion peptide, heptad repeat one, and the central helix domains, which are likely subject to greater functional constraints (Fig. 4A).

**Fig. 4.**
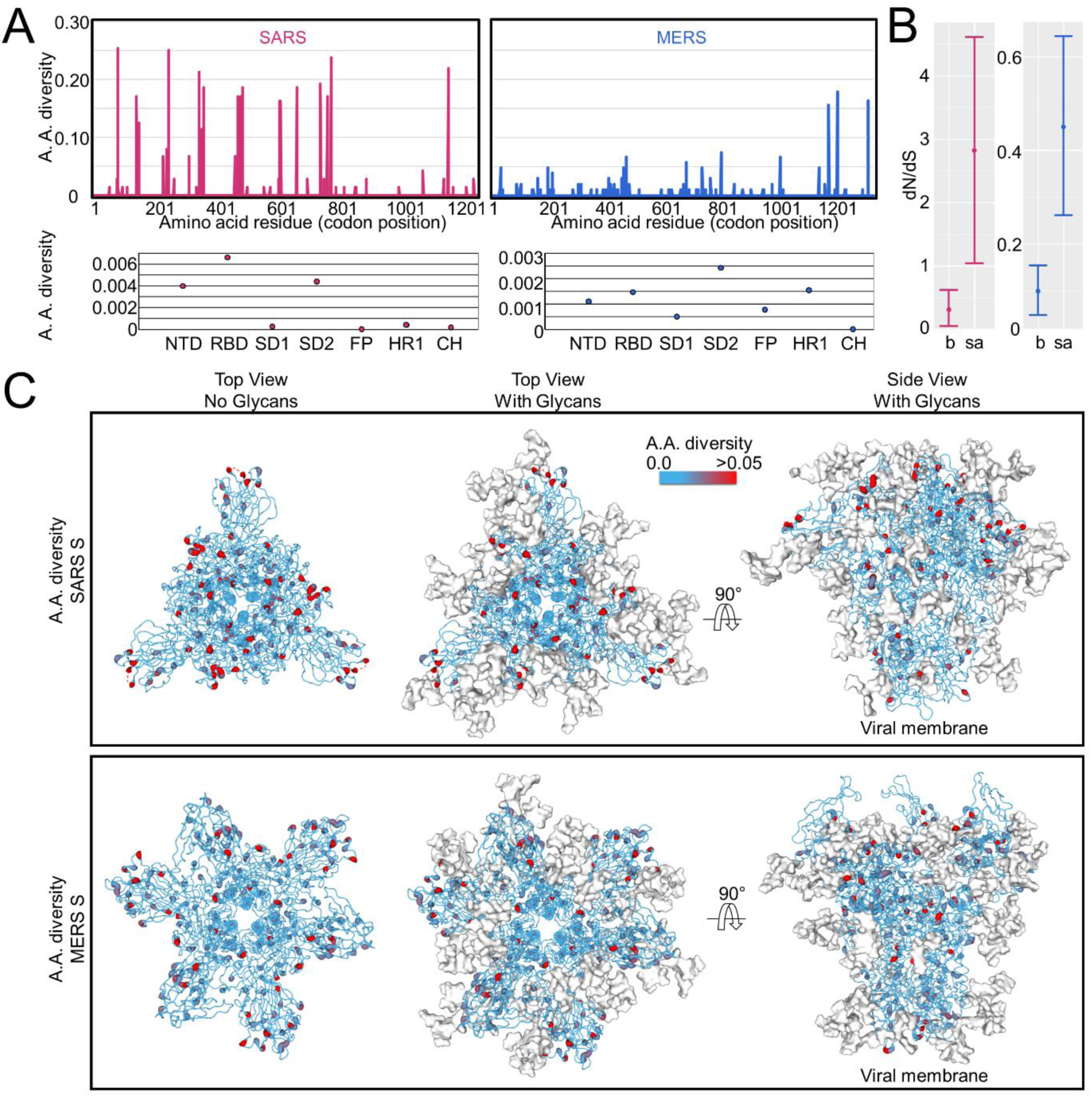
Amino-acid sequence diversification across SARS and MERS spikes. **(A)** Amino-acid diversity in SARS and MERS S gene sequences. Averaged values for each domain are also shown below. **(B)** Comparison of dN/dS values between buried and exposed residues across SARS and MERS S. The error bars correspond to the 95% highest posterior density intervals while the circles indicate mean dN/dS values. **(C)** Mapping of the per residue amino-acid diversity shown in **A** onto the structures of SARS and MERS S (PDB ID 5X58 and 5X59, respectively)^17^. S proteins are presented as backbone traces with residues colored according to amino-acid diversity. Residues with elevated diversity are colored in red, and N-linked glycans are presented as white surfaces.

Analysis of the relative ratio of non-synonymous to synonymous nucleotide substitutions (i.e. dN/dS ratios) revealed that exposed residues exhibited significantly higher dN/dS values (Fig. 4B). Buried residues on SARS had mean dN/dS ratios of 0.31 compared to 2.82 for exposed resides. Likewise, the buried residues on MERS had a calculated dN/dS ratio of 0.10 compared to exposed residues with a value of 0.45. Furthermore, when per-site amino-acid diversities were mapped onto the fully glycosylated structural model of the respective CoV S proteins (Fig. 4C), hotspots of mutations were highlighted on the protein surface throughout the trimer revealing extensive vulnerabilities permeating through the glycan shield of SARS and MERS CoVs.

Although dN/dS estimates are comparable within each viral outbreak, they are not directly comparable between viral families as they can only be considered in the environment in which they are measured (i.e. multiple differences in transmission ecology and host-virus interactions disallow meaningful comparisons). For example, differences in the epidemic behaviour and host immune environment of MERS and SARS outbreaks likely contribute to the observed genetic diversity and thus dN/dS. MERS was characterized by repeated spillover events from camels into humans, where it circulated transiently. In contrast, the SARS outbreak corresponded to a single zoonotic event followed by extensive human-to-human transmission. Consequently, inferring the degree of selection acting upon MERS and SARS from dN/dS analysis is extremely difficult.

### Visualisation of HIV-1 Env and SARS S glycan shields by cryo-EM analysis

HIV-1 Env is a prototypic viral class I fusion protein that exhibits extensive surface glycosylation, resulting in an effective glycan shield to aid evasion from the host adaptive immune response^25,52^. In order to visualize the structure of the respective glycan “shields” of HIV-1 and SARS coronavirus, we used single-particle cryo-electron microscopy (cryo-EM), as recently described^53^. The results for HIV-1 Env were reproduced from Berndsen et al.^53^ while the previously published SARS 2P dataset^54^ was reprocessed for this study. Although cryo-EM datasets of fully glycosylated MERS S^41^ and chimpanzee simian immunodeficiency virus (SIVcpz)^55^ are also available, only the HIV and SARS data were of sufficient quality (Fig. 5). Using combination of low-pass filtering and thresholding, along with 3D variability analysis, we can reveal the previously hidden structure of the SARS glycan shield and compare it with the HIV-1 Env glycan shield^53^ (Fig. 5). We observe the nearly all-encompassing glycan density on HIV-1 Env and evidence for extensive glycan-glycan interactions, especially in the oligomannose patch regions, whereas the glycans on SARS S are more isolated and lack the wide-ranging glycan networks that are the hallmark of an effective glycan shield^56,57^. The high variability around the S1 receptor binding domains is indicative of their flexibility, which is necessary for the spike to engage its receptor^58^; however, this intrinsic property complicates the analysis of the signal from glycans in this region.

**Fig. 5.**
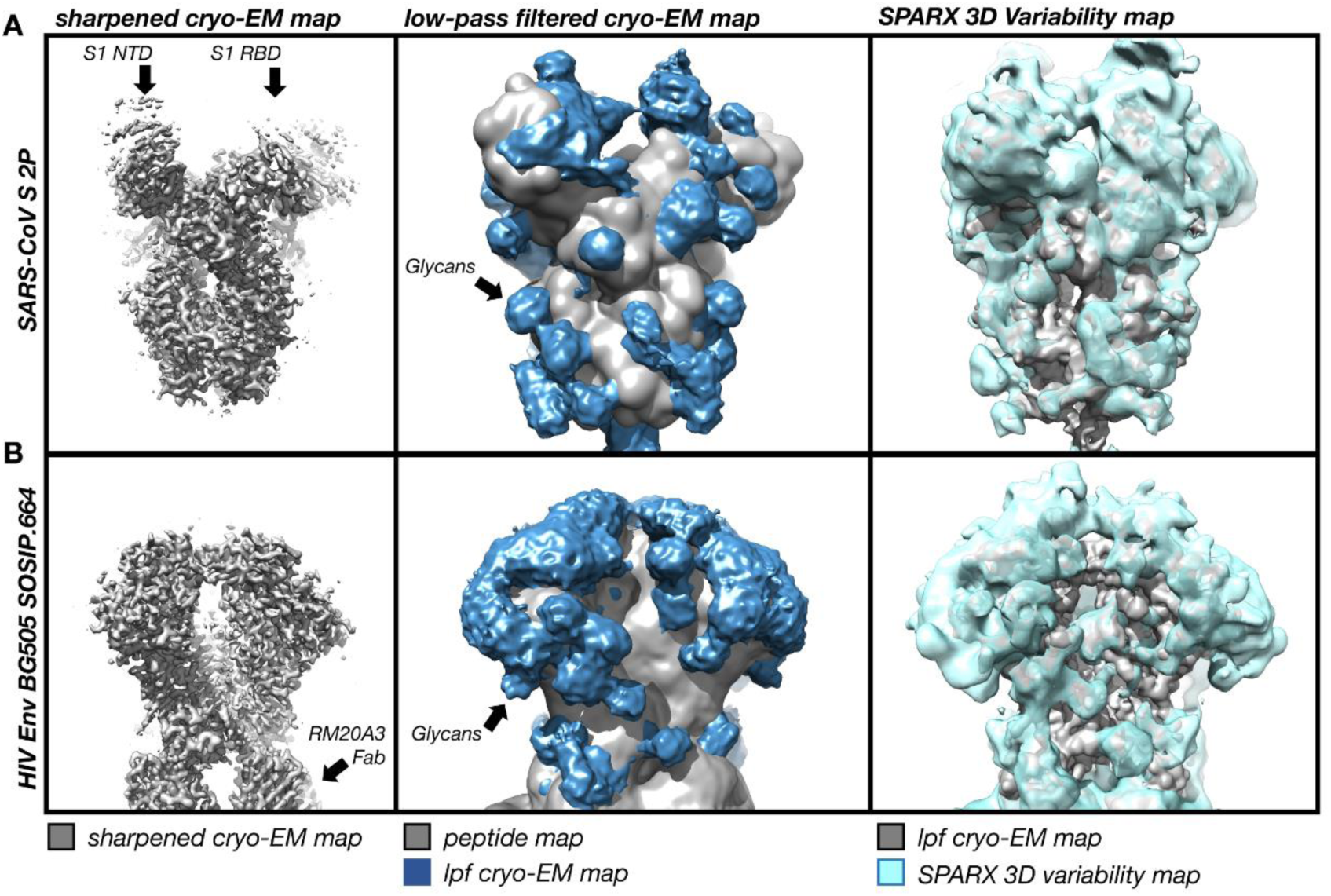
Comparative cryo-EM analysis of SARS S and HIV-1 Env glycan shields. **(A)** Left panel: Sharpened 3.2Å-resolution C3-symmetric cryo-EM map of SARS S 2P ectodomain^54^ visualized at a high contour level with disordered S1 receptor binding and N-terminal domains extending out from the central core. Middle panel: Low-pass filtered cryo-EM map of the glycoprotein visualised at a low contour level along with a simulated peptide-only map overlaid. Right panel: SPARX 3D variability map^53^. **(B)** Same as in **(A)** but for HIV-1 Env BG505 SOSIP.664 construct in complex with 3 copies of RM20A3 base-specific Fabs^53^.

### Comparison of Viral Glycosylation Reveals Disparate Shielding Efficacies

Viral envelope proteins are glycosylated to varying degrees, but depending on their overall mass, surface area, and volume, the overall density of glycan shielding may differ significantly. For example, both LASV GPC and coronavirus S proteins consist of 25% glycan by molecular weight. However, given the significantly larger protein surface area and volume of coronavirus S proteins, coverage of the glycan “shield” over the proteinaceous surface is considerably sparser in comparison to the smaller LASV GPC, which occludes a far greater proportion of the protein surface with fewer glycans. To demonstrate that the presence of glycosylation plays a major role in the immune response to these different glycoproteins, we studied the glycome of several biomedically important coronaviruses and compared their glycan compositions in a structural context.

We then investigated the glycan shield densities of seven viral class I fusion proteins using a global structural approach which was calculated by dividing the number of amino-acids that interact with glycans by the number of solvent-accessible amino-acid residues of each respective glycoprotein and plotted this against oligomannose abundance. A strong correlation was observed (Fig. 6) and viruses historically classified as “evasion strong”^59^ had significantly elevated glycan shield densities and oligomannose abundance, which underscores the importance of glycan shielding in immune evasion.

**Fig. 6.**
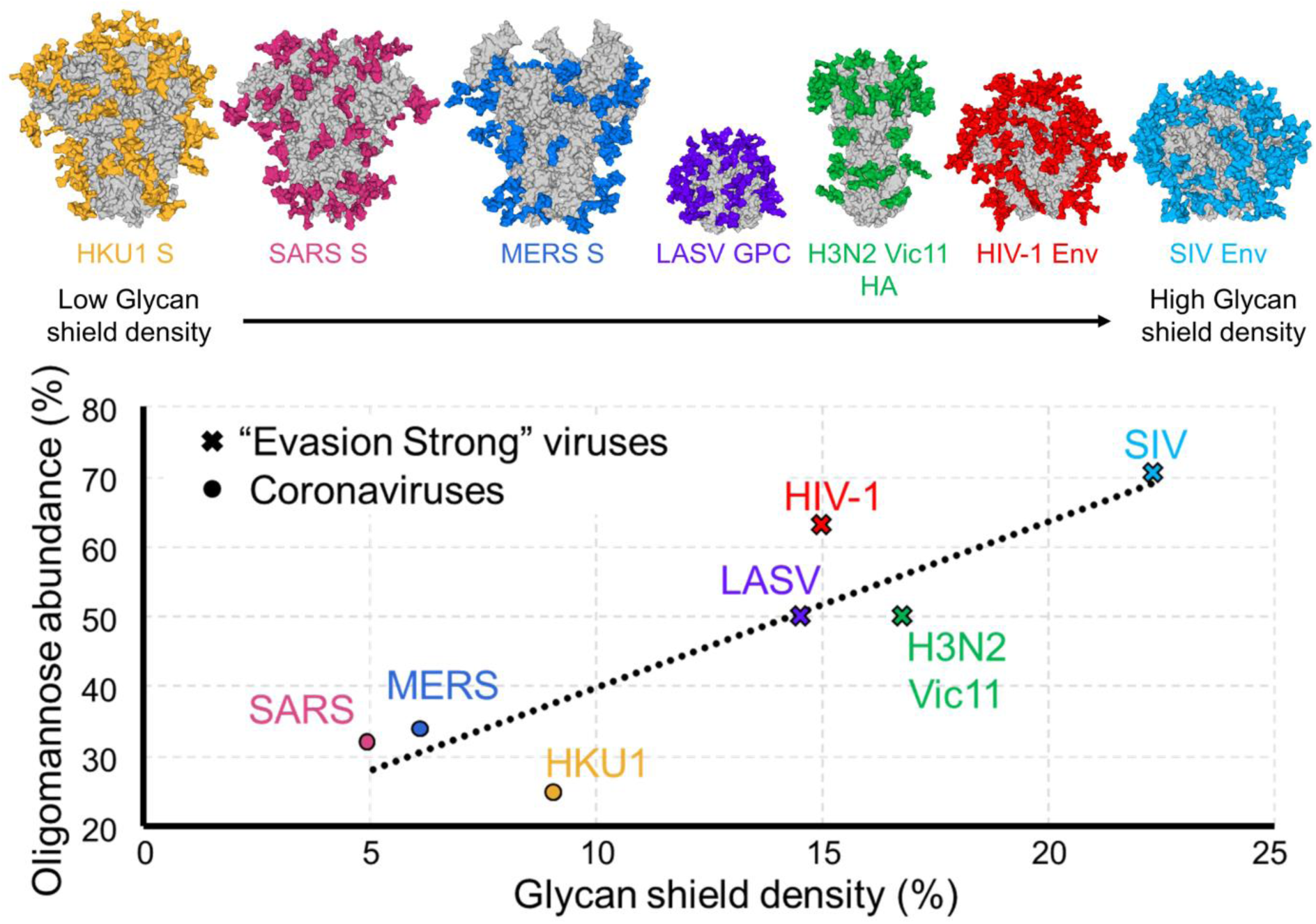
Comparison of the glycan shields of viral class I fusion proteins. Glycan shield densities were calculated using Proteins, Interfaces, Structures and Assemblies (PISA)^60^ analyses of fully glycosylated models of SARS S, MERS S, HKU1 S, LASV GPC, HIV-1 Env (BG505), Influenza H3N2 hemagglutinin (Victoria 2011), SIV Env (PDB ID 5X58, 5X59, 5I08, 5VK2, 4ZMJ, 4O5N, 6OHY, respectively)^15,17,55,61–63^. Oligomannose abundances of viral glycoproteins were ascertained by HILIC UPLC analysis of PNGase F released N-linked glycans that were fluorescently labelled with procainamide^29,46,55^ (SI Fig. 5). The number of amino-acid residues interacting with N-linked glycans was divided by the number of solvent-accessible amino-acid residues of the glycoprotein as a measure for global glycan shield density. All viral glycoproteins analysed were expressed as trimers in HEK 293F cells apart from LASV GPC, which was derived from virus-like particles from Madin-Darby canine kidney II cells.

Whether the restricted glycan shielding observed on coronaviruses is linked to the zoonosis of the pathogens is unknown. However, it is tempting to speculate, for example, that MERS has not evolved a dense shield since it would not offer as much of a protective advantage against camel nanobodies (also known as single-domain antibodies) which could more easily penetrate it. Investigation of the host immune response to viruses in their natural reservoirs may offer a route to understanding why coronavirus glycosylation does not reach the density of other viruses such as HIV-1. Additionally, it may be that functional constraints, such as maintaining flexibility of the receptor binding domains, limit the accretion of glycans on coronavirus spikes, which would render it incapable of performing its primary functions, including receptor binding and membrane fusion. This phenomenon has been observed on other viral glycoproteins, including influenza HAs, where there is a limit to the accumulation of glycosylation sites that can be incorporated *in vivo*^64,65^, compared to *in vitro*^66^, with H3N2 and H1N1 HAs replacing existing PNGs rather than continually adding them upon the glycoprotein^26,65^.

More topically, it is interesting to note the conservation of N-linked glycosylation sites on S proteins from the novel coronavirus found in the Wuhan in 2019 and SARS (SI Fig. 6). The Wuhan 2019-nCoV possesses a total of 22 N-linked glycan sites compared to 23 on SARS, with 18 of these sites being in common. As such, it is likely that these glycans on this novel coronavirus would shield similar immunogenic epitopes that are observed on SARS S. As expected, most of the differences between the two viruses are observed on the S1 subunit, due to its amenability to substitutions whilst still remaining functionally competent. Furthermore, likely targets for the majority of antibodies targeting the spike are located on S1, resulting in greater levels of immune pressure upon this subunit. This notion is further reflected in terms of glycosylation, with all of the glycan sites conserved on the S2 subunit between SARS and 2019-nCoV, whereas the S1 subunit exhibits glycan site additions and deletions (SI Fig. 7).

Although it is difficult to directly compare viruses in terms of immunogenic responses, on the one hand, SARS and MERS coronaviruses readily elicit neutralizing antibodies following infection or immunization^67–70^. Indeed, many potential MERS CoV vaccine candidates are able to elicit high titres of serum IgG upon immunization but fail to produce sufficient mucosal immunity^70^. In contrast, the high mutation rate^71^ and the evolving glycan shield of HIV-1^24^, which firmly exemplifies it as “evasion strong” virus, hinders the development of broadly neutralizing antibodies^72^.Viruses classified as “evasion strong”^31,59^ may then differ due to varied efficacies of protein surface shielding by glycans.

Overall, this study reveals how the extensive N-linked glycan modifications of SARS and MERS CoV S proteins do not constitute an effective shield, which is reflected by the overall structure, density and oligomannose abundances across the trimeric glycoproteins. We also demonstrate that amino-acid diversification indeed occurs at antibody accessible regions on the trimer, which confirms that glycans play a role in occluding specific regions if vulnerability on the glycoprotein. Furthermore, comparisons between glycan shields from a number of viruses highlight the importance of a glycan shield in immune evasion and reveal structural principles that govern glycosylation status.

## Methods

### Expression and purification of coronavirus spike glycoproteins

Human embryonic kidney 293 Freestyle (HEK293F) cells were transfected with mammalian-codon-optimised genes encoding 2P-stabilised SARS MERS and HKU1 S proteins, as previously described ^41^. H3N2 Victoria 2011 hemagglutinin was also expressed in the HEK 293F cells. Cultures were harvested 6 days after transfection, filtered and purified by nickel-affinity chromatography and size exclusion chromatography using a Superdex™ 16/600 75 pg column (GE Healthcare).

### Release and labelling of N-linked glycans

Excised coronavirus S gel bands were washed alternately with acetonitrile and water before drying in a vacuum centrifuge. The bands were rehydrated with 100 μL of water and incubated with PNGase F at 37 °C overnight. Aliquots of released N-linked glycans were also fluorescently labelled with procainamide, by adding 100 μL of labelling mixture (110 mg/mL procainamide and 60 mg/mL sodium cyanoborohydrate in 70% DMSO and 30% glacial acetic acid) and incubating for 4h at 65 °C. Labelled glycans were purified using Spe-ed Amide 2 columns (Applied Separations), as previously described ^33^.

### Glycan analysis by HILIC UPLC

Labelled glycans were analysed using a 2.1 mm x 10 mm Acquity BEH Glycan column (Waters) on an Acquity H-Class UPLC instrument (Waters), as performed previously ^33^, with fluorescence measurements occurring at λ_ex_ = 310 nm and λ_em_ = 370 nm. Quantification of oligomannose-type glycans was achieved by digestion of fluorescently labelled glycans with Endo H, and clean-up using a PVDF protein-binding membrane (Millipore). Empower 3 software (Waters) was used for data processing.

### Mass spectrometry of glycans

Ion-mobility electrospray ionisation MS and tandem MS of released N-linked glycans were performed on a Synapt G2Si instrument (Waters) as previously described ^46^. N-linked glycans were purified on a Nafion^®^ 117 membrane (Sigma-Aldrich) prior to injection. Data acquisition and processing were carried out using MassLynx v4.11 and Driftscope version 2.8 software (Waters).

### Mass spectrometry of glycopeptides

Aliquots of 30-50 μg of coronavirus spikes were denatured, reduced and alkylated as described previously^36^. Proteins were proteolytically digested with trypsin (Promega), chymotrypsin (Promega), alpha-lytic protease (Sigma-Aldrich) and Glu-C (Promega). Reaction mixtures were dried and peptides/glycopeptides were extracted using C18 Zip-tip (MerckMilipore) following the manufacturer’s protocol. Samples were resuspended in 0.1% formic acid prior to analysis by liquid chromatography-mass spectrometry using an Easy-nLC 1200 system coupled to an Orbitrap Fusion mass spectrometer (Thermo Fisher Scientific). Glycopeptides were separated using an EasySpray PepMap RSLC C18 column (75 μm x 75 cm) with a 240-minute linear solvent gradient of 0-32% acetonitrile in 0.1% formic acid, followed by 35 minutes of 80% acetonitrile in 0.1% formic acid. Other settings include an LC flow rate of 200 nL/min, spray voltage of 2.8 kV, capillary temperature of 275 °C, and an HCD collision energy of 50%. Glycopeptide fragmentation data were extracted form raw files using Byonic™ (Version 3.5.0) and Byologic™ (Version 3.5-15; Protein Metrics Inc.). Glycopeptide fragmentation data were manually evaluated with true-positive assignments given when correct b- and y- fragments and oxonium ions corresponding to the peptide and glycan, respectively, were observed. The extracted ion chromatographic areas for each true-positive glycopeptide, with the same amino-acid sequence, were compared to determine the relative quantitation of glycoforms at each specific N-linked glycan site.

### Model construction

Structural models of N-linked glycan presentation on SARS, MERS and HKU1 S were created using electron microscopy structures (PDB ID 5X58, 5X59, and 5I08, respectively)^15,17^, along with complex-, hybrid-, and oligomannose-type N-linked glycans (PDB ID 4BYH, 4B7I, and 2WAH). The most dominant glycoform presented at each site was modelled on to the N-linked carbohydrate attachment sites in Coot^73^.

### Molecular evolution analysis

Publicly available sequences encoding full-length GPC spike gene for SARS-CoV (3765bp) were downloaded from GenBank and manually aligned. For MERS-CoV, we leveraged the whole genome alignment collated by Dudas et al.^74^.Specifically, the alignment corresponding to the spike gene was extracted (4059bp), excluding sequences isolated from humans. Final alignments for SARS- and MERS-CoV corresponded to 70 and 100 sequences, respectively.

For the dN/dS analysis, we first estimated Bayesian molecular clock phylogenies for SARS- and MERS-CoV independently using BEAST v 1.8.4^75^. For both viruses, we assumed an uncorrelated log-normal distributed molecular clock^76^, Bayesian Skyline coalescent prior^77^ and a codon-structured substitution model^78^ Multiple independent MCMC runs of 10-20 million steps were executed to ensure stationarity and convergence had been achieved. Empirical distributions of time-scaled phylogenies were obtained by combining (after the removal of burnin) the posterior tree distributions from the separate runs, which were subsequently used to estimate dN/dS ratios using the renaissance counting approach ^79,80^ implemented in BEAST v 1.8.4. We also estimated per-site amino-acid diversity, which was calculated as the average number of amino-acid difference between two sequences at an amino-acid position in all possible pairs in the sequence alignment.

### Cryo-EM data analysis and visualization

Single-particle cryo-EM data analysis of BG505 SOSIP.664 in complex with RM20A3 Fab was reproduced from Berndsen et al.^53^. Data for the SARS-CoV S 2P ectodomain was previously published^54^ and reprocessed for this analysis according to the methods outlined by Berndsen et al.^53^. In summary, both datasets were acquired on a FEI Titan Krios (Thermo Fisher) operating at 300 KeV equipped with a K2 Summit Direct Electron Detector (Gatan). Movie micrographs were aligned and dose weighted with MotionCor2^81^ and CTF estimation was performed with Gctf^82^. Single-particle data processing was performed using CryoSparc v.2^83^ and Relion v.3^84^. 3D variability analyses were performed in SPARX^85,86^. Map filtering and visualisation was performed in UCSF chimera^87^.

### Clustering analysis of viral glycan shields

Interactions between N-linked glycans and amino-acid residues were calculated using Proteins, Interfaces, Structures and Assemblies (PISA) European Bioinformatics Institute (EBI)^60^. Glycan shield density was calculated by the number of amino-acid residues interacting with glycans divided by the total number of amino-acid residues.

## Acknowledgements

The Wellcome Centre for Human Genetics is supported by grant 203141/Z/16/Z. We thank the Medical Research Council (MR/S007555/1 to T.A.B.), NIH (R56 AI127371 to I.A.W., R01 AI127521 to J.S.M. and A.B.W.), Bill and Melinda Gates Foundation (grants OPP1115782 to A.B.W., and OPP1170236 to I.A.W. and A.B.W.), the International AIDS Vaccine Initiative, Bill and Melinda Gates Foundation through the Collaboration for AIDS Discovery (grants OPP1084519 to M.C., and 1196345 to I.A.W., A.W.B., and M.C.), and the Scripps Consortium for HIV Vaccine Development (CHAVD) (AI144462 to M.C., A.B.W., and I.A.W.).

