## Supplementary material for "Vulnerabilities in coronavirus glycan shields despite extensive glycosylation": Supplmentary Information

This document includes Supplementary Figures 1-7, Supplementary Figure Legends, and Supplementary References.

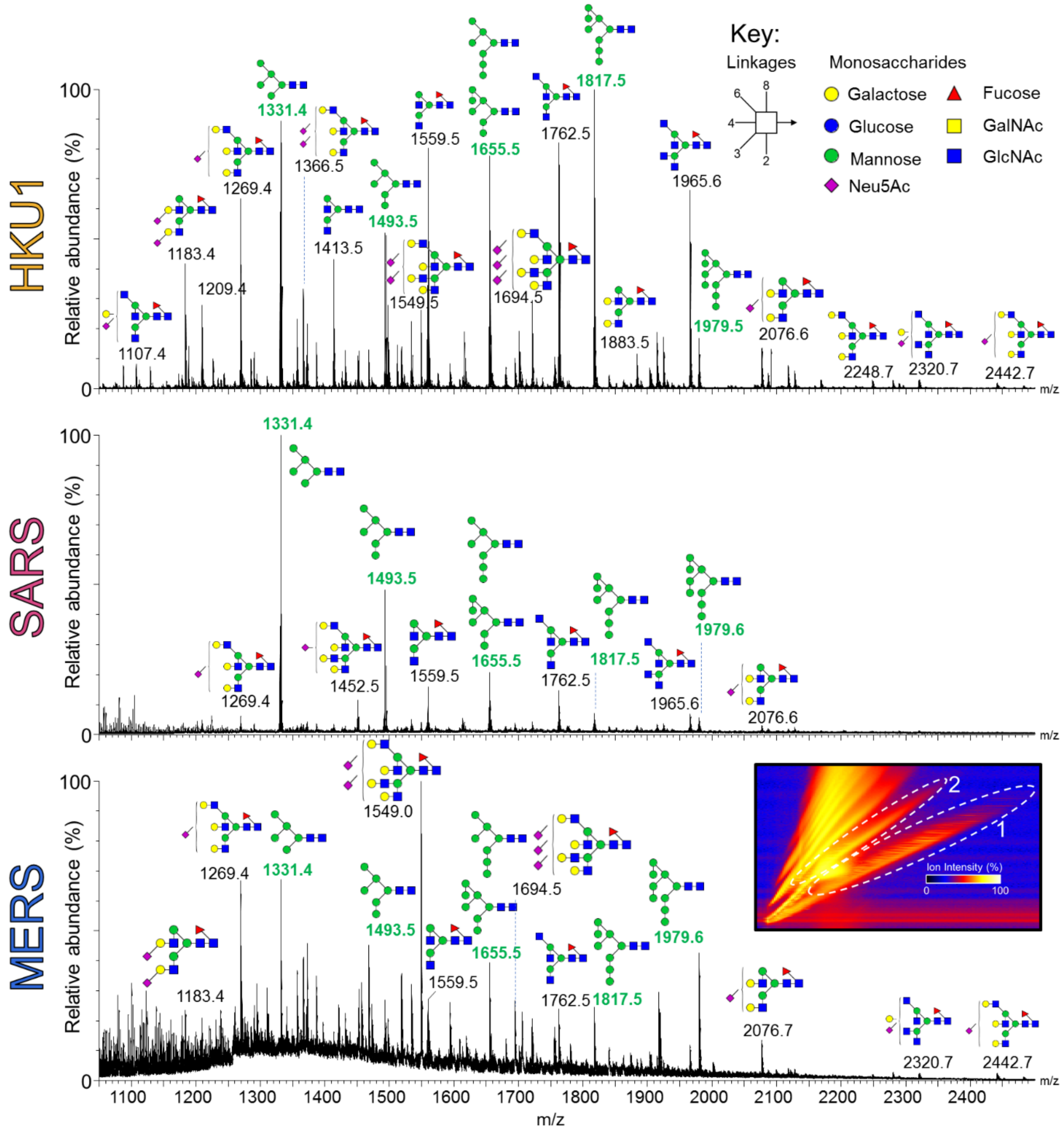


**Fig. S1.** Ion mobility-extracted mass spectra of singly- and doubly- charged N-linked glycan ions from HKU1, SARS and MERS S glycoproteins. Peaks are annotated with the corresponding compositions, using Consortium for Functional Glycomics symbolic nomenclature and Oxford system linkages[1], as per the key. Oligomannose-type glycan *m/z* values are labelled in green.

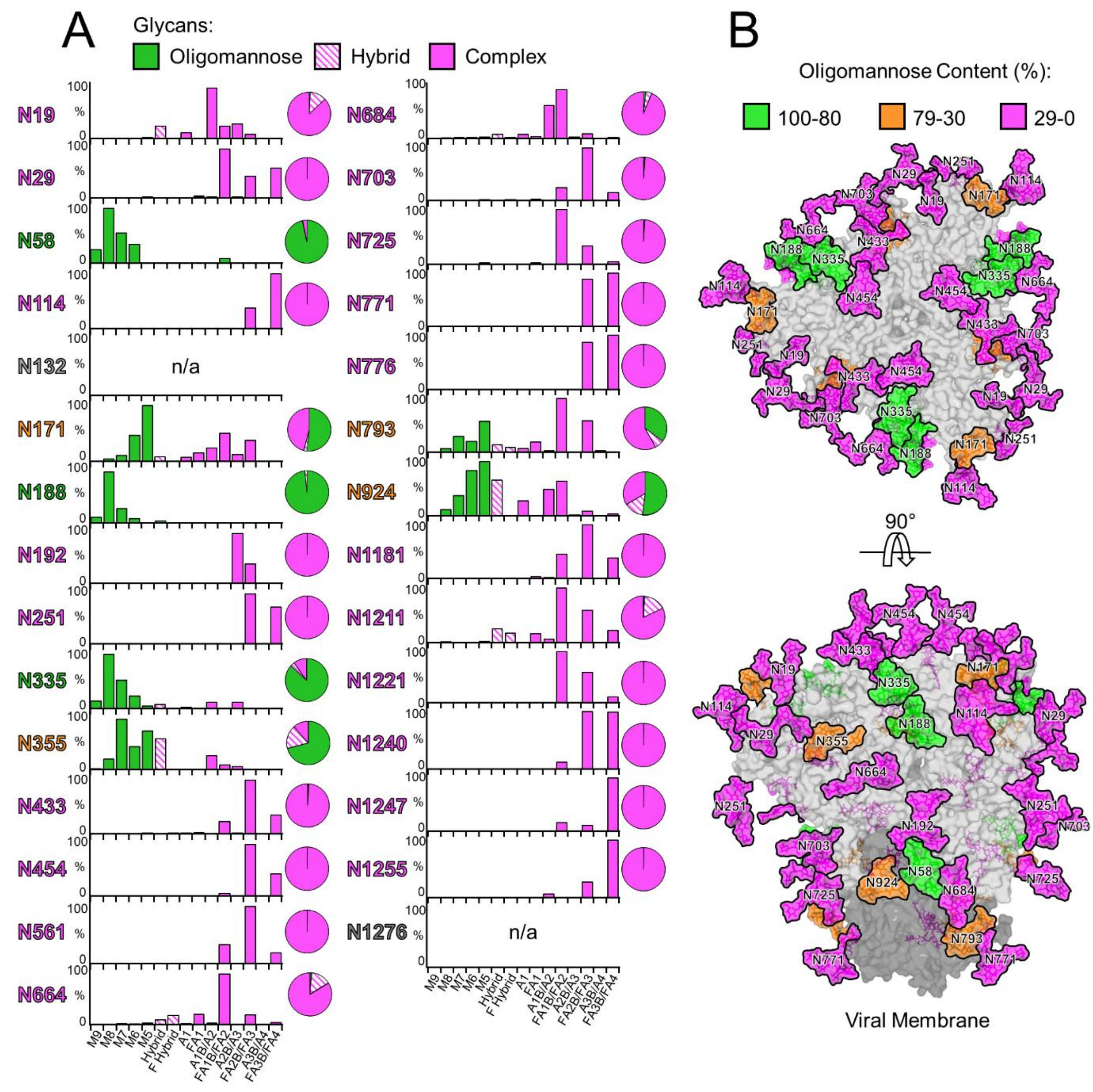


**Fig. S2.** Compositional analysis and structure-based mapping of HKU1 S N-linked glycans. (A) Quantitative site-specific N-linked analysis of HKU1 S. Purified HKU1 S was digested with trypsin, chymotrypsin, and trypsin + chymotrypsin, then analysed by LC-ESI MS. Glycan compositions are based on the glycan library generated from negative-ion mass spectrometry of released N-glycans. The bar graphs represent the relative quantities of each glycan group with oligomannose-type glycan series (M9 to M5; $Man_9GlcNAc_2$ to $Man_5GlcNAc_2$) (green), afucosylated and fucosylated hybrid glycans (Hybrid & F Hybrid) (dashed pink), and complex glycans grouped according to the number of antennae and fucosylation (A1 to FA4) (pink). Left to right; least-processed to most processed. The pie charts summarise the quantification of these glycans. (B) Modelling of experimentally observed glycosylation onto the pre-fusion structure of trimeric HKU1 S (PDB ID code 5I08)[2]. The glycans are coloured according to oligomannose content, as defined by the upper right-hand key. S1 and S2 subunits coloured light grey and dark grey, respectively.

| MERS | N66 | N104 | N125 | N155 | N166 | N222 | N236 | N244 | N410 | N487 | N592 | N619 | N719 | N774 | N785 | N870 | N1176 | N1213 | N1225 | N1241 | N1256 | N1277 | N1288 |
|---|---|---|---|---|---|---|---|---|---|---|---|---|---|---|---|---|---|---|---|---|---|---|---|
| Fucosylation (%) | 0 | 0 | 0 | 0 | 0 | 0 | 3 | 0 | 0 | 100 | 100 | 100 | 100 | 99 | 99 | 26 | 100 | 19 | 100 | 100 | 100 | 100 | 0 |
| Sialylation(%) | 0 | 18 | 0 | 0 | 0 | 0 | 0 | 28 | 0 | 25 | 38 | 22 | 30 | 31 | 27 | 19 | 66 | 0 | 0 | 7 | 0 | 0 | 0 |

| SARS | N29 | N65 | N73 | N109 | N118 | N119 | N158 | N227 | N269 | N318 | N330 | N357 | N589 | N602 | N691 | N699 | N783 | N1056 | N1080 | N1116 | N1140 | N1155 | N1176 |
|---|---|---|---|---|---|---|---|---|---|---|---|---|---|---|---|---|---|---|---|---|---|---|---|
| Fucosylation (%) | 97 | 0 | 91 | 98 | 99 | 99 | 0 | 0 | 75 | 98 | 97 | 77 | 0 | 64 | 98 | 51 | 5 | 83 | 12 | 100 | 96 | 100 | 97 |
| Sialylation(%) | 18 | 11 | 17 | 22 | 54 | 69 | 0 | 0 | 22 | 32 | 28 | 68 | 0 | 14 | 3 | 99 | 1 | 16 | 36 | 0 | 4 | 0 | 35 |

| HKU1 | N19 | N29 | N58 | N114 | N132 | N171 | N188 | N192 | N251 | N335 | N355 | N433 | N454 | N561 | N664 | N684 | N703 | N725 | N771 | N776 | N793 | N924 | N1181 | N1211 | N1221 | N1240 | N1255 | N1276 |
|---|---|---|---|---|---|---|---|---|---|---|---|---|---|---|---|---|---|---|---|---|---|---|---|---|---|---|---|---|
| Fucosylation (%) | 17 | 100 | 3 | 100 | n/a | 33 | 0 | 48 | 100 | 0 | 2 | 99 | 100 | 100 | 92 | 57 | 99 | 99 | 100 | 100 | 58 | 16 | 99 | 87 | 100 | 100 | 96 | n/a |
| Sialylation(%) | 8 | 16 | 0 | 0 | n/a | 0 | 0 | 48 | 0 | 0 | 0 | 8 | 0 | 13 | 29 | 31 | 11 | 23 | 0 | 0 | 3 | 1 | 1 | 8 | 53 | 0 | 0 | n/a |

**SI Fig 3.** Site-specific quantification of fucosylation and sialylation of N-linked glycan sites, on MERS, SARS, and HKU1 S glycoproteins.

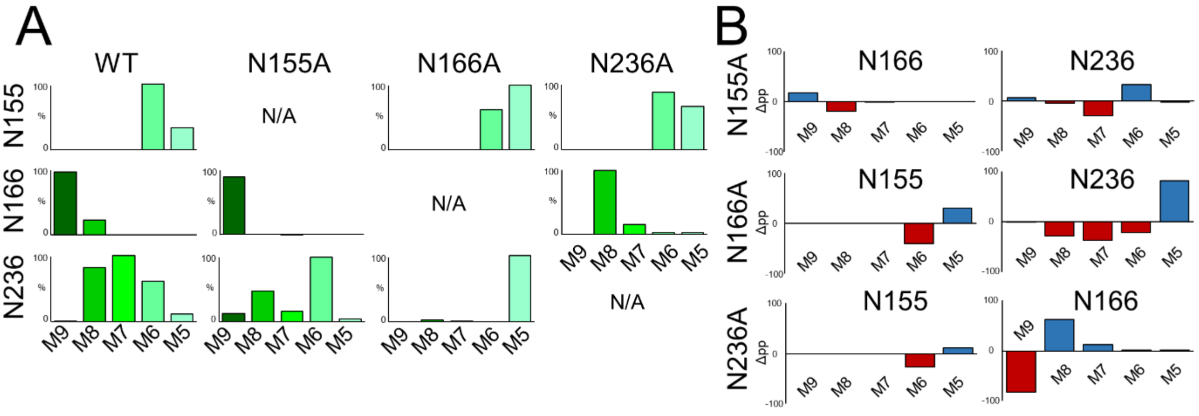


**SI Fig 4.** Glycan deletion increases mannose trimming of N-linked glycans on MERS S oligomannose patch. (A) Relative quantitation of glycans in the mannose patch sites (N155, N166, N236) on MERS S. M9 to M5; ($Man_9GlcNAc_2$ to $Man_5GlcNAc_2$) (Dark green to Pale green). (B) Percentage point differences in the abundance of oligomannose-type glycans at mannose patch sites in the glycan knockout mutants compared to WT MERS. Decreased and increased abundances are colored red and blue, respectively.

| | MERS S | SARS S | HKU1 S | LASV GPC[3] | HIV-1 BG505 Env[4] | SIV MT145K Env[5] | H3N2 Vic11 HA |
|---|---|---|---|---|---|---|---|
| Oligomannose-type (%) | 33.8 | 32.2 | 25.0 | 49.5 | 63.0 | 70.5 | 50.1 |
| Complex-type (%) | 66.2 | 67.8 | 75.0 | 50.5 | 37.0 | 29.5 | 49.9 |

**SI Fig. 5** Oligomannose- and complex-type glycan composition table of viral fusion proteins, quantified by HILIC-UPLC. Endo H digestions of labelled glycans were performed to measure oligomannose abundance.

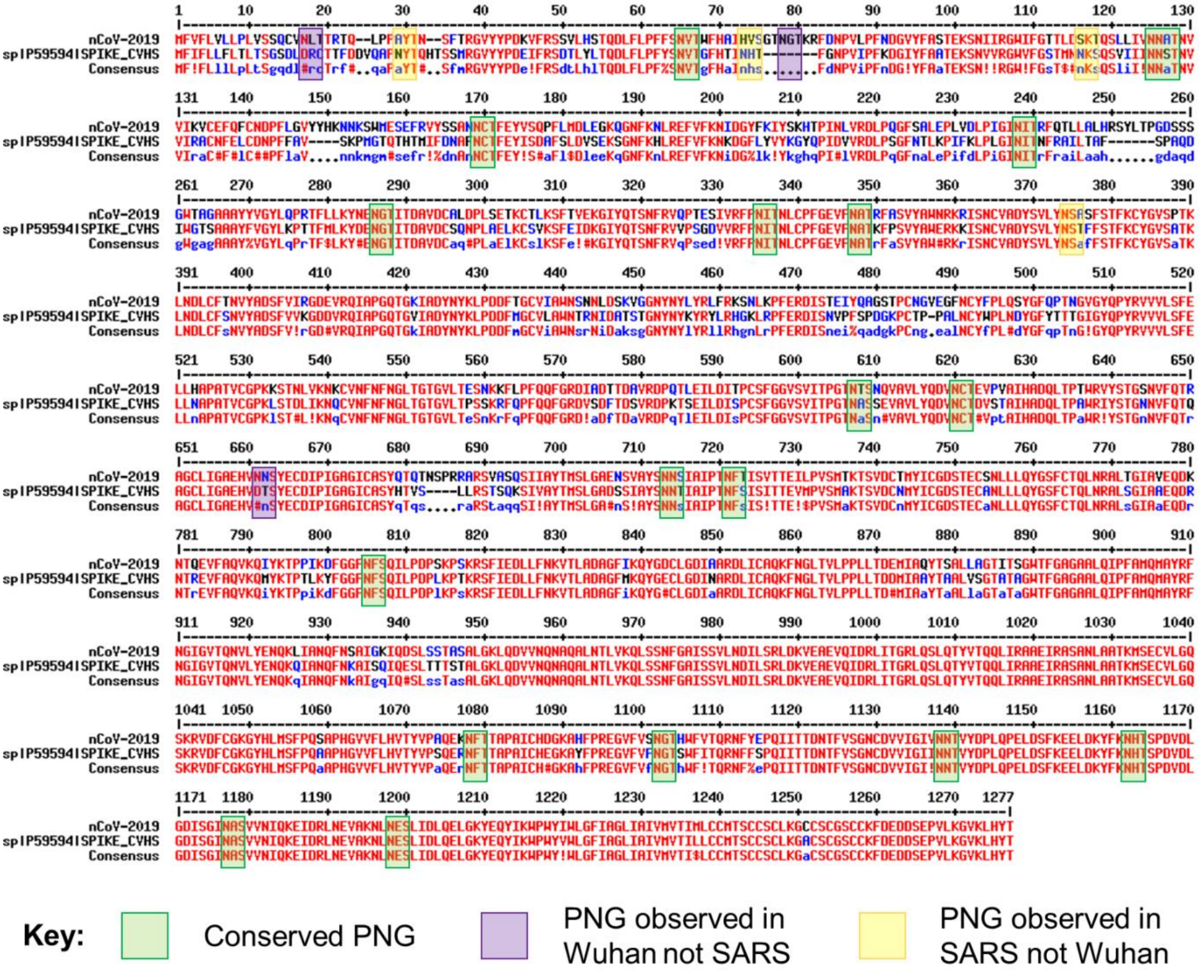

**SI Fig. 6:** Sequence alignment of S proteins nCoV-2019 from Wuhan (Genbank: MN908947.3) and SARS CoV (Uniprot: P59594) highlighting conservation of N-linked glycan sequons. Conserved potential N-linked glycosylation sites (PNGs) are colored in green, PNGs observed in nCoV-2019 but not in SARS are colored purple, and PNGs observed in SARS but not nCoV-2019 are colored in yellow.

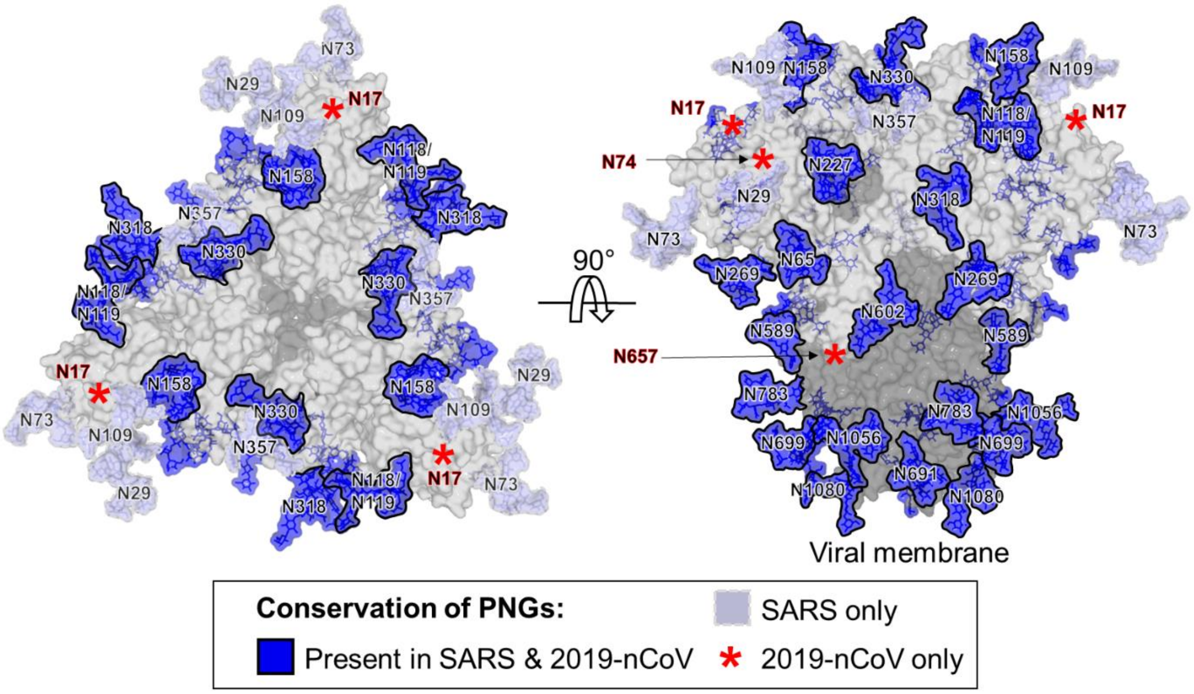


**SI Fig 7**: Mapping the conservation of glycosylation site between SARS and 2019-nCoV. Glycan sites were modelled onto SARS S (PDB ID:5X58)[6], with glycan sites conserved between both viruses, colored blue. Glycan sites only present in SARS are colored in light blue-grey. Approximate positions of N-linkned glycans present on 2019-nCoV are highlighted by red asterisks, with numbering based on the 2019-nCoV protein sequence.
